# Qudaich: A smart sequence aligner

**DOI:** 10.1101/060509

**Authors:** Sajia Akhter, Robert A Edwards

## Abstract

Next generation sequencing (NGS) technology produces massive amounts of data in a reasonable time and low cost. Analyzing and annotating these data requires sequence alignments to compare them with genes, proteins and genomes in different databases. Sequence alignment is the first step in metagenomics analysis, and pairwise comparisons of sequence reads provide a measure of similarity between environments. Most of the current aligners focus on aligning NGS datasets against long reference sequences rather than comparing between datasets. As the number of metagenomes and other genomic data increases each year, there is a demand for more sophisticated, faster sequence alignment algorithms. Here, we introduce a novel sequence aligner, Qudaich, which can efficiently process large volumes of data and is suited to de novo comparisons of next generation reads datasets. Qudaich can handle both DNA and protein sequences and attempts to provide the best possible alignment for each query sequence. Qudaich can produce more useful alignments quicker than other contemporary alignment algorithms.

**Author Summary:** The recent developments in sequencing technology provides high throughput sequencing data and have resulted in large volumes of genomic and metagenomic data available in public databases. Sequence alignment is an important step for annotating these data. Many sequence aligners have been developed in last few years for efficient analysis of these data, however most of them are only able to align DNA sequences and mainly focus on aligning NGS data against long reference genomes. Therefore, in this study we have designed a new sequence aligner, qudaich, which can generate pairwise local sequence alignment (at both the DNA and protein level) between two NGS datasets and can efficiently handle the large volume of NGS datasets. In qudaich, we introduce a unique sequence alignment algorithm, which outperforms the traditional approaches. Qudaich not only takes less time to execute, but also finds more useful alignments than contemporary aligners.

## Introduction

Many sequence alignment tools have been developed in last couple of decades. There are sensitive local alignment tools like BLAST (Altschul et al., 1990) and FASTA (Pearson & Lipman, 1988) that were developed in the 1990’s. A number of methods for faster sequence alignments like SSAHA2 (Ning, Cox & Mullikin, 2001), BLAT (Kent, 2002), MUMmer (Delcher et al., 1999, Delcher et al., 2002; Kurtz et al., 2004) have been published since 2000. Several new, fast and memory efficient algorithms have been developed since 2008 because of the next-generation sequencing technologies that generate millions of short sequence reads. These tools include BWA (Li & Durbin, 2009, Li & Durbin, 2010), Bowtie (Langmead & Salzberg, 2012), Maq (Li, Ruan & Durbin, 2008), Soap (Li et al., 2008; Liu et al., 2012) etc. Most of the sequence aligners developed in the last few years mainly concentrate on aligning next generation sequencing datasets against a long reference genome. None of the recently developed aligners focus on generating alignments between the datasets created by shorter read sequencing technologies. For pair wise comparisons between any two datasets produced by NGS, general-purpose alignment tools like USEARCH (Edgar, 2010) or BLAST (Altschul et al., 1990) are more effective. However, the total time required for these comparisons is still a great concern. Some of the fast sequence aligners (like Bowtie, SOAPaligner, BWA-short), which align short reads against long reference sequences might be useful for this purpose, but it is not guaranteed that those aligners can efficiently handle a database containing hundreds of thousands or millions of reads instead of a few long reference sequences. There are many applications that require comparison of (typically) short read data sets with each other. For example, in metagenomic analyses pairwise comparisons between datasets provide a rapid assessment of the similarity between sites (Cassman et al., 2012; Dutilh et al., 2012), allow the identification of novel genome sequences (Dutilh et al., 2014; Brown et al., 2015), and provide estimates of organism abundance to characterize different environments (Lim et al.).

Another concern of current alignment algorithms is that most of the fast sequence alignment approaches developed in last few years mainly produce DNA sequence alignments, but protein sequence alignments are also essential. Very few alignment algorithms (e.g. BLAST, Usearch, MUMmer, RAPSearch) support protein sequence alignments or translated nucleotide sequence alignments. Therefore, for protein sequence alignments, there are still opportunities for improved alignments algorithms that can provide protein sequence

In this paper, we present a new sequence aligner, Qudaich (**q**ueries and **u**nique **d**atabase **a**lignment **i**nferred by **c**lustering **h**omologs) that introduces a novel approach to the sequence alignment problem. The main design purpose of qudaich is to focus on datasets from next generation sequencing, i.e. to find the alignment between two different NGS datasets, instead of aligning the NGS query dataset against a long reference sequence. The NGS datasets generally have hundreds of thousand sequences or more, and so the input database for qudaich could contain large number of sequences. Qudaich is flexible and its algorithmic structure imposes no restriction on the absolute limit of the acceptable read length, but the current version supports read lengths less than 2,000 base pairs. Qudaich can be used to align both DNA and protein sequences.

## Methods

Qudaich (available on https://github.com/linsalrob/qudaich/) performs local sequence alignments in two major steps - i) identifying the candidate database sequence(s) and ii) generating the optimal alignment with those candidate database sequences.

In the first step, qudaich tries to find the candidate database sequence(s) for each query sequence. Here, for a query sequence, *q*, the candidate database sequence refers to the corresponding database sequence, *d* that gives either the best alignment or very close to the best alignment with *q*. Thus, if *q* is aligned against all the database sequences, the alignment score between *q* and *d* will be either the best score or very close to the best score. Qudaich mainly focuses on short-read datasets and such databases typically contain huge numbers of sequences, therefore any naïve approaches to find the candidate database sequences are impractical. Qudaich applies heuristics to limit the search space to find the candidate database sequences efficiently. After identifying all the candidate database sequences, the second step is to generate the optimal alignment for each query sequence with the corresponding candidate database sequence. The Smith-Waterman-Gotoh algorithm (SWG) (Gotoh, 1982) is used for this purpose.

The approach taken by qudaich has a key advantage over most of the contemporary aligners that are based on suffix tree or hash based approaches. These aligners normally consider a seed (a match of some length) to generate the alignment, but suffer from the problem that the optimal alignment may not contain that particular seed. Qudaich, on the other hand, does not apply heuristics to generate the alignments. It only applies heuristics to find the candidate database sequences.

### Identifying the candidate database sequence

Qudaich uses a novel algorithmic structure to look for the candidate database sequences efficiently. Most aligners keep the database and query sequences separate, but in qudaich both database and query sequences are organized together. This novel organization has two main advantages. It accelerates the searching of candidate database sequences by clustering several query sequences; and, it allows us to construct powerful heuristics to limit the search space. However, the primary disadvantage of using the query and database sequences together is that the sequence indices need to be rebuilt for each different comparison.

### Primary data structure

Qudaich uses a suffix array as the primary data structure. It constructs a single suffix array using all database and query sequences. Among the several approaches or data structures used in the sequence alignment algorithms described earlier, the suffix array is one of the most efficient ways to store or organize the sequences. In 2008, Ge Nong *et al.* (Nong, Zhang & Chan, 2011) provided two efficient algorithms for constructing suffix arrays and later, Yuta Mori (“sais”) provided a faster implementation of the suffix array construction library based on those algorithms. This implementation is both time and space efficient – it runs in *O(n)* time and requires *MAX(2n, 4k)* extra working space where *n* is the length of the string and *k* is the size of the alphabet in the string. We adopted this library to construct the suffix array.

### Suffix array organization in qudaich

In our algorithm, a long string, *S* is constructed by concatenating all the database sequences, the reverse complement of the database sequences and then all the query sequences. Then using *S*, a suffix array, *SA* is constructed.

There can be two types of suffixes in the suffix array, because all the database sequences and query sequences are concatenated in *S*. Some suffixes start with either a query sequence or with a part of a query sequence. These are called *Qry* suffixes. The rest of the suffixes start with either database sequences or with a part of the database sequences or with the reverse complement of the database sequence. These suffixes are called DB suffixes. In the suffix array, all consecutive *Qry* suffixes construct a *Qry* group. Similarly, all consecutive *DB* suffixes make a *DB* group, where each group contains one or more suffixes. As shown in the Figure 1, the suffix array, *SA* contains a sequence of *Qry* groups and *DB* groups. So, *SA* will be a sequence of alternating *Qry* groups and *DB* groups.

**Figure 1.**
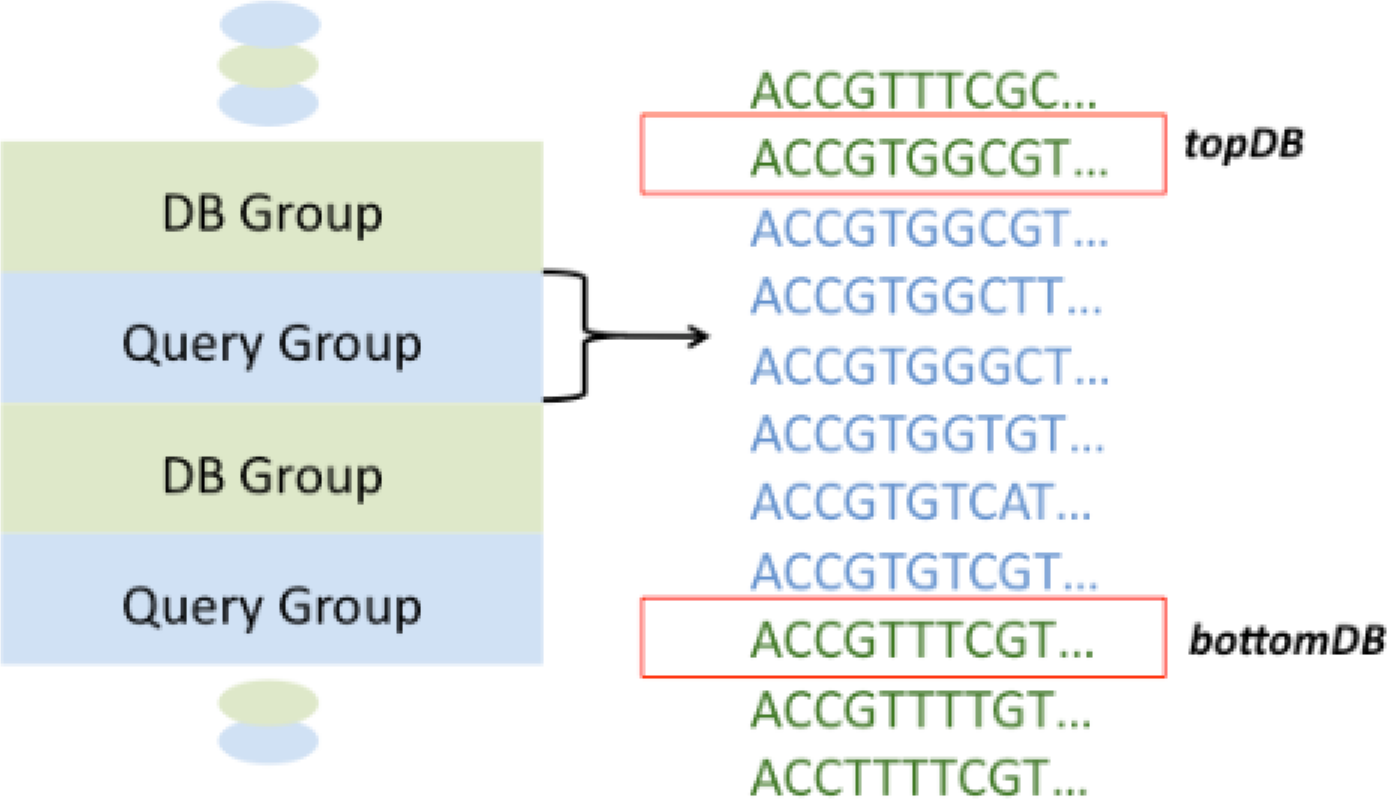
Order of the database suffixes and query suffixes in the suffix array. The red boxes show the two database suffixes, one of which has the longest match with the query suffixes that are marked as light blue.

### Algorithm Overview

All the suffixes in the suffix array are stored in sorted order. Based on this characteristic and the arrangement of *Qry* groups and *DB* groups, it is possible to find the longest exact match for all the query suffixes with the database suffixes.

Let us assume a *Qry* group occurs in position *SA[i]* to *SA[j]* in the suffix array *SA* where i ≤ j. The best match of all the *Qry* suffixes of this group will be the immediately prior *DB* suffix (*topDB*; Figure 1), *SA[i−1]* and/or the immediately subsequent *DB* suffix (*bottomDB*; Figure 1), *SA[j*+*1]* of the group, because all the suffixes in *SA* are in lexicographically sorted order. Therefore, only two *DB* suffixes need to be considered for each group of query suffixes instead of all *DB* suffixes. This statement holds for all the *Qry* groups in the suffix array. So the search space is limited to all the *Qry* groups and their surrounding two *DB* suffixes. Note that, in the suffix array construction, the reverse complement of the database sequences is used instead of using the reverse complement of query sequences to minimize the number of *Qry* groups and reduce the processing time.

The suffix array data organization in qudaich finds the best exact match for each query suffix. This property can be further exploited to construct two different heuristics that can be used to find the candidate database sequences (Figure 2).

**Figure 2.**
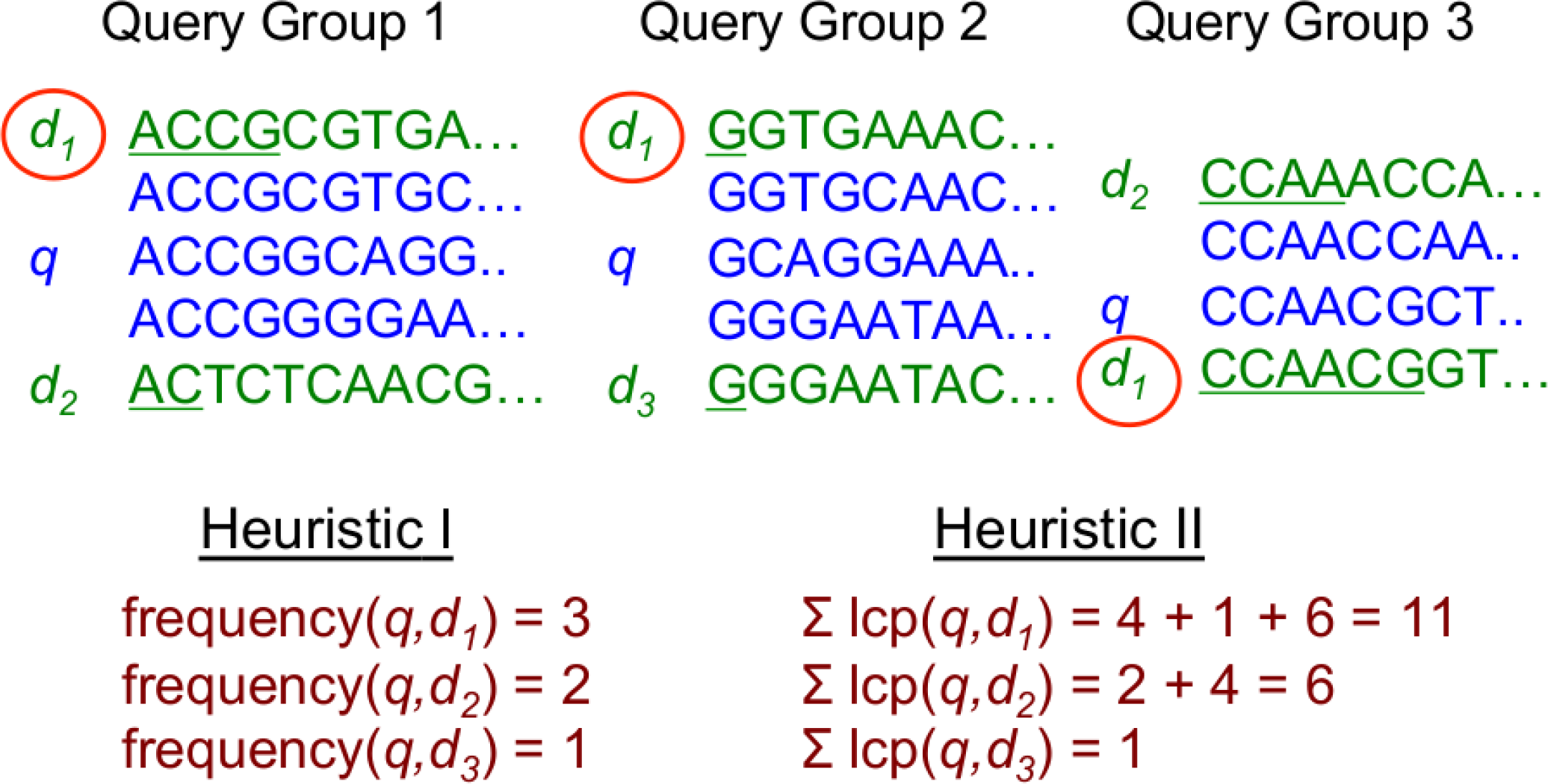
Example of the selection of candidate database sequence for query *q* using heuristic I and heuristic II. Both heuristics select *d_1_* as the candidate database sequence for *q*.

*Heuristic I: Query q has the best alignment with database sequence d, if the suffixes of d are the most frequent closest suffixes in all the query groups containing all the suffixes of q*.

This heuristic is validated by the following statement: if *q* has the best alignment with *d*, the number of common prefix matches (of length at least one) between the suffixes of *q* and the corresponding suffixes of *d* will be the maximum among the number of the common prefix match between the suffixes of *q* and the suffixes of any other database sequence. Since all the suffixes are lexicographically sorted in *SA*, in most of the cases suffixes of *d* (that have some match with suffixes of *q*) will be either *topDB* or *bottomDB* of the *Qry* groups containing the corresponding suffixes of *q*.

*Heuristic II: Query q has the best alignment with database sequence d, if heuristic Σ satisfies and L lcp(suffixes of q, suffixes of d) is maximal*.

For heuristic II, a weighted frequency is used. Instead of only counting the presence of the *DB* suffix in *topDB* and *bottomDB*, the longest common prefix (lcp) between the query suffix and the *DB* suffix is also measured.

The heuristics described above mainly use the topDB and bottomDB to find the candidate database sequences. However, this might not give the best accuracy in some circumstances. For convenience, we demonstrate the concept assuming that there is a query sequence *q* and we have the longest exact matches for each suffix of *q* (that is with tobDB and bottomDB).

As shown in Figure 3, occasionally a suffix of q, say *SA[k]* in a *Qry* group from *SA[i]* to *SA[j]* where *i* ≤ *k* ≤ *j*, has the longest match with topDB at *SA[i−1]* and the match length is *a*. But *SA[i−2]* (which is a DB suffix) also has a match up to length *a* with *SA[k]a*. However, the database sequence (say d_1_), whose suffix is at *SA[i−2]*, has a better alignment with *q*. Since *SA[i−2]* was not considered in heuristic I or II, d_1_ might not be predicted as a candidate database sequence. In the ideal case, more than one DB suffix in the *DB* group needs to be considered for the all query suffixes in the corresponding query group. Hence, another possible heuristic is discussed here, which incorporates both heuristic I and heuristic II.

**Figure 3.**
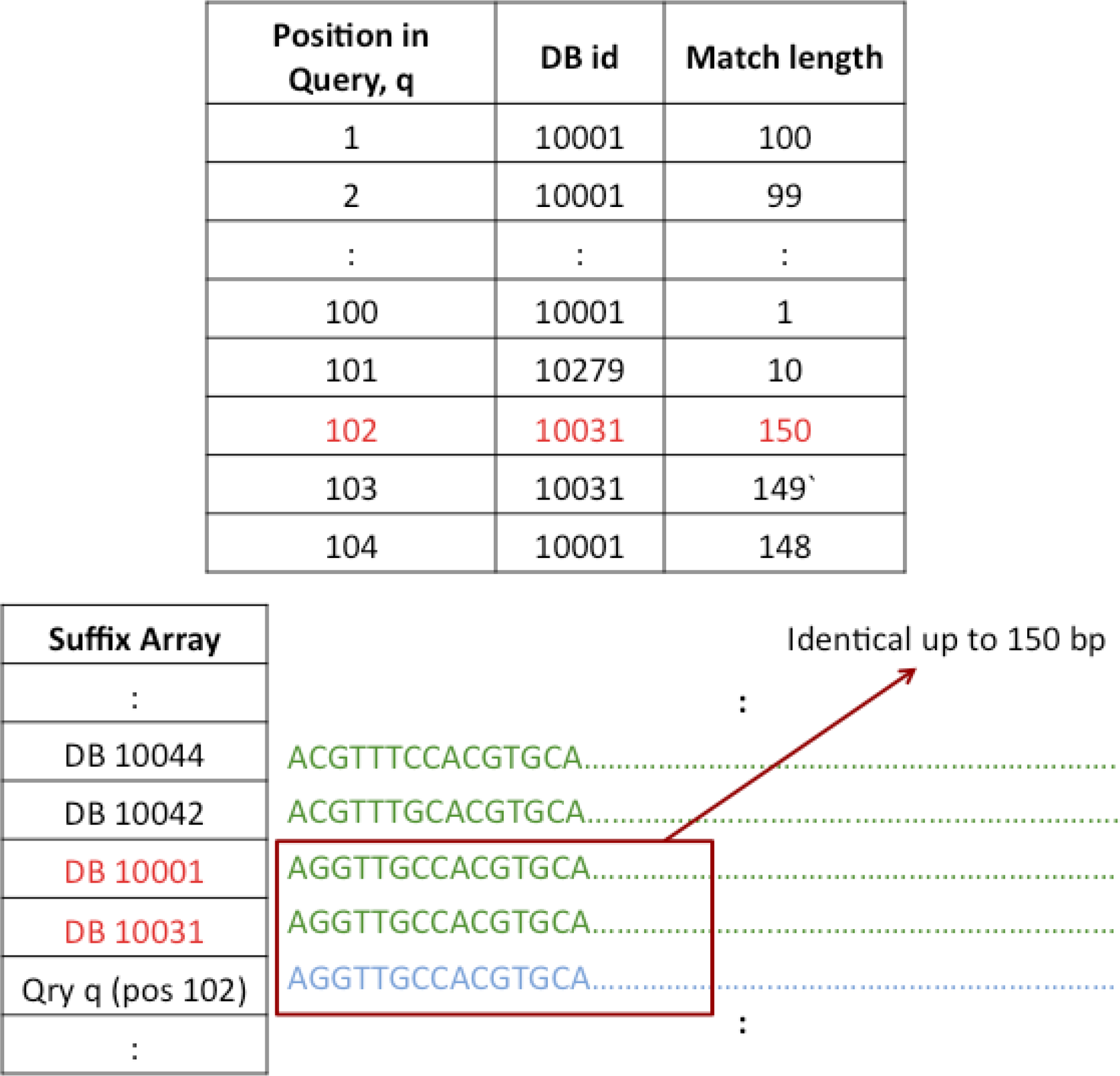
The top table shows the information of each query suffix of query *q* after finding the longest exact matches and the bottom table shows the ambiguity order of the suffix array. The query *q* has a match with DB 10001 from position 1 to 100 and there is a mismatch at position 101 and then there is a match from position 102 to 151. The position 102 and position 103 shows that suffixes start from these positions have match with DB 10031 instead of DB 10001. This happens because the suffix that matches at position 102 of query *q*, is identical up to 150 bp with both DB 10031 and DB 10001.

*Possible heuristic III:* In this heuristic, more than one DB suffix will be considered while calculating heuristic I and heuristic II. For a query group *SA[i]* to *SA[j]*, where i≤ j, both heuristics I and II consider only topDB at *SA[i−1]* and bottomDB at *SA[j+1]*. An alternative approach is to consider multiple DB suffixes from both top and bottom DB group of the query group. That means instead of just considering topDB at *SA[i−1]* and bottomDB at *SA[j+1], SA[i−1]* to *SA[i−1−x]* will be considered as topDBs and similarly, *SA[j+1]* to *SA[j+1+x]* will be considered as bottomDBs.

Initial empirical testing has suggested that the absolute best hit is always within the top five sequences (and is usually the adjacent sequence), and thus the additional overhead and significant increases in runtime for this calculation does not appear to benefit the algorithm. However, if it is possible to design an ideal heuristic, which picks the candidate database sequences that always gives the best alignment, qudaich will always generate a 100% accurate alignment. This is the key benefit of the algorithmic approach of qudaich.

In qudaich, heuristics I and II were implemented to find the candidate database sequences. The implementation details of these heuristics are discussed below.

### Implementation of identifying the candidate database sequences

Two different procedures have been followed for the implementation of heuristic I and heuristic II.

To find the candidate database sequences using heuristic I, the suffix array needs to be traversed only once and the following steps taken.

i. For all *Qry* suffixes of each query group, store the corresponding *topDB* and *bottomDB*.
ii. For each query sequence, sort the frequency of the database sequences.
iii. Use the most frequent database sequence as the candidate database sequence for each query sequence.

To explore the complexity of this approach, suppose there are *r* query sequences of length *n* and *x* different database suffixes, which appear as *topDB* or *bottomDB*. This procedure requires *O(r. xlog(x))* time. Since *x* will be less than or equal to *2n* as there are *n* suffixes for each query sequence and for each query suffix two *DB* suffixes are stored. In the worst case, when there is no good alignment for a query sequence, *x* will be close to or equal to *2n*. Otherwise, *x* should be small. Thus in the worst case, the time complexity will be *O(r.n)* while in the best case it will approach *O(r)* (i.e. the number of query sequences). The space requirement will be *O(r.n)*.

For heuristic II, the longest common prefix (lcp) between two suffixes needs to be calculated. A lcp array holds the lcp information of two consecutive suffixes. That means, *lcp[i]* holds the length of the longest common prefix between *SA[i]* and *SA[i−1], lcp[i+1]* contains the lcp between *SA[i]* and *SA[i+1]* and so on, where 1 ≤ *i* ≤ length of *SA*. It is possible to construct the lcp array while constructing the suffix array. Using Yota Mori’s suffix array implementation, Johannes Fischer efficiently computed the lcp array. This implementation outperforms other lcp array construction algorithms. Therefore, this library was adopted for heuristic II (“elventear/sais-lite-lcp”).

The lcp array only provides the lcp between two consecutive suffixes. But the lcp between all the suffixes in each query group and their *topDB* and *bottomDB* need to be calculated. Therefore, a dynamic programming approach that is linear in terms of time complexity was designed for this purpose. The lcp between a query suffix at position *i* and a *DB* suffix is computed using the following recurrence relation:

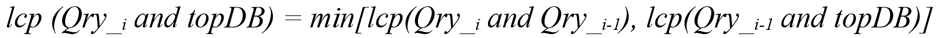

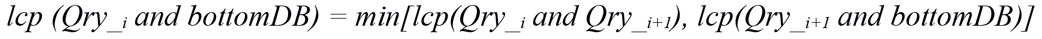

Suppose there is a query group from *SA[i]* to *SA[j]* where *i*<*j*. The topDB is at *SA[i−1]* and bottomDB is at *SA[j+1]*. We need to calculate the lcp between *SA[i−1]* (topDB) and query suffixes from *SA[i]* to *SA[j]*. From *lcp[i]*, we get the lcp between *SA[i−1]* and *SA[i]*, say *lcp[i]* = *x*. *lcp[i+1]* holds the lcp between *SA[i+1]* and *SA[i]* (say *lcp[i+1]* = *y*). The lcp between *SA[i−1]* and *SA[i+1]* will be the minimum between *x* and *y*, because the suffix at *SA[i−1]* matches up to *x* length with the suffix at *SA[i]* and the suffix at *SA[i]* matches up to *y* length with the suffix at *SA[i+1]*. As all the suffixes are lexicographically sorted in *SA*, when *x*< *y*, the suffix at *SA[i+1]* has to match up to *x* length with the suffix at *SA[i−1]* and vice versa when *y*>*x*. Assume the lcp between *topDB* and *SA[i+1]* is *z*. To find the lcp between *topDB* and *SA[i+2]*, the minimum between *z* and *lcp[i+2]* will be calculated based on the above fact. So by calculating the lcp between *SA[i]* and topDB, we can calculate the lcp between *SA[i+1]* and topDB and so on. The same procedure is followed for calculating the lcp between bottomDB and query suffixes from *SA[i]* to *SA[j]* (Figure 4).

**Figure 4.**
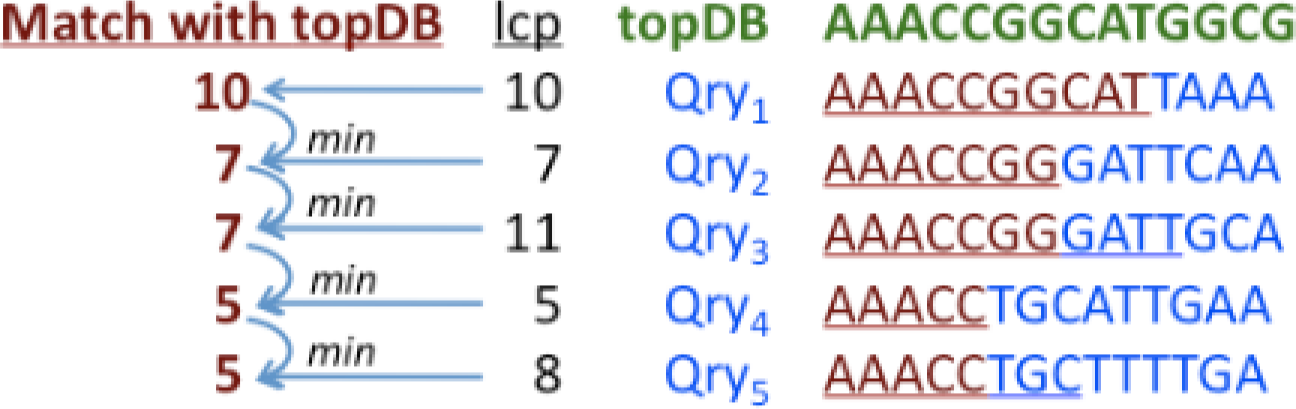
Calculation of the longest common prefix (lcp) between *topDB* and *Qry* suffixes from *Qry_1_* to *Qry_5_*. The lcp between *Qry*_1_: and *topDB* is obtained from the lcp array and that is 10. The lcp between Qry_2_ and *topDB* is the minimum value from lcp(*Qry_1_*, *Qry_2_*) and match(*Qry_1_* and *topDB*), which is 7. Similarly, the lcp between *Qry3* and *topDB*, *Qry_4_* and *topDB*, and *Qry_5_* and *topDB* are 7, 5 and 5 accordingly.

### Generating the alignment

For each query sequence, the candidate database sequence(s) is identified in the previous step. Next, the Smith-Waterman-Gotoh algorithm (Gotoh, 1982) is applied to generate the optimal alignment between each query sequence and the candidate database sequence using the user defined scoring matrix and gap penalties. In general, the Smith-Waterman-Gotoh algorithm produces the optimal alignment with affine gap penalties and the time complexity is quadratic. Suppose there are *m* query sequences of length *r* and *n* database sequences of length *l*. In our case, this algorithm will not be costly, because the time complexity will be *O(mrl)* instead of *O(mrnl)* where *r* and *l* will typically be < 2000 bp and *m* and *n* will be in excess of 10^4^ and often exceeds 10^6^.

### Options to run Qudaich

Like other aligners, qudaich has several options that can be defined by the users to get better alignments based on different datasets. In qudaich, while the candidate database sequences are identified for each query sequence, it is possible to find more than one candidate database sequence per query sequence based on the heuristic that is used. Qudaich reports the average frequency (heuristic I) or the average lcp length (heuristic II) of all the candidate database sequences for all query sequences. By default, qudaich generates alignments for only those query sequences where the lcp length or frequency is greater than the average of those. Users can also choose different values for lcp length or frequency cutoffs (using the same option -a) to produce alignments greater than that input value. Therefore, if the database and the query dataset have no significant matches, the average frequency or lcp length value should be very low and user can use a larger value (than the average) for the option -a. Also, if the database and the query dataset are very similar and the user wants more than one alignment per query sequence, they can first generate more candidate database sequences per query sequence and then use a smaller value (than the average) for the option-a to generate the alignments. The alignment score is the score generated by the Smith-Waterman-Gotoh algorithm that uses user defined match/mismatch score and gap opening/extending penalties.

## Results

Qudaich generates alignments based on the Smith-Waterman-Gotoh algorithm that has already been shown to produce the optimal alignment (Gotoh, 1982). Therefore, the accuracy of the alignment test is unnecessary, and the main concern is to prove how accurately the candidate database sequences are selected.

### Testing datasets

Three different test sets were used to test the accuracy and the speed of the aligner.

Test set 1: For a detailed comparison between several aligners, both the database and the query dataset were created from *Escherichia coli O157:H7 str. EDL933* (5,528,445 bp). Grinder (Angly et al., 2012) was used to create shotgun and amplicon sequence libraries from this genome. There were 10,000 sequences of 100 bp in both database and query dataset.

Test set 2: Two published mosquito metagenomes were used as the first test set. The first mosquito metagenome with 32,965,046 base pairs was used as the database and the other mosquito metagenome with 54,339,014 base pairs was used as the query dataset. The database contained 318,477 sequences with an average sequence length of 103 basepairs. The average sequence length of query dataset was also similar, 101 basepairs and the dataset contained 533,720 sequences (Dinsdale et al., 2008).

Test set 3: The third test set was designed to test protein sequence alignments. Genemark (Zhu, Lomsadze & Borodovsky, 2010) was used to identify open reading frames from the sequences in test set 2, and these were converted to amino acid sequences. The query mosquito metagenome contains 343,611 sequences of length 31 aa (on average) and the database contains 177,418 sequences of average length 32 aa.

### Accuracy for finding candidate database sequences

A detailed comparison between several aligners was performed using test dataset 1 to test the accuracy of the algorithms. We compared qudaich to the blastn program from BLAST+, USEARCH, YASS and EMBOSS WATER (Rice, Longden & Bleasby, 2000; Noé & Kucherov, 2005; Camacho et al., 2009; Edgar, 2010) (the commands used to run these programs are shown in the appendix). First, for each pair of programs, we show the number of query sequences where both algorithms agree with each other. That means for query *q*, if alignment algorithm A and alignment algorithm B both identify the same candidate database sequence *d*, we say that A and B agree that *q* has a best match with database sequence *d*. As shown in table 1 (highlighted in blue), blastn and qudaich agree for most query sequences (7,470 queries using heuristic I; 8,172 queries using heuristic II). However, YASS (6,525 queries), WATER (3,466 queries) and USEARCH (6,576 queries) also agree more with blastn than with other algorithms. For qudaich, the candidate database sequences that do not have a match with blast generally have poor alignment scores (Figure 5). Second, if alignment algorithm A and alignment algorithm B agree for the alignment between query *q* and database sequence *d*, we consider it a *valid alignment* if and only if the location of *q* and the location of *d* overlaps in the genome (note that there might be a better alignment between *q* and another database sequence *d_1_* where *q* and *d_1_* do not overlap). For the *valid alignment* too, blastn and qudaich agree more than any other pair of alignment algorithms (Table 1, highlighted in green). Also, blastn produces most alignments (for 4,665 queries) where the query sequences have overlaps with the corresponding database sequences. Qudaich has the second most alignments (4,603 queries using heuristic I; 4,625 queries using heuristic II) where the query sequences and the candidate database sequences overlap. In summary, blastn has the most alignments in common with other algorithms, in contrast to WATER that has the least alignments in common with other algorithms. WATER uses the Smith-Waterman algorithm to calculate the optimal local alignment between two sequences. However, the optimal alignment between two sequences does not necessarily infer biological similarity between the sequences unless the edit distance between both sequences is very small. Qudaich produces an optimal alignment between the query and candidate database sequences because it uses heuristics to find the candidate sequences before the alignment step.

**Table 1:** Comparison between qudaich and several other aligners for finding the accuracy of the candidate database sequences using dataset 1. Blue cells show the number of query sequences in common between two alignment approaches where both aligners find the same candidate database sequence for the corresponding query sequence. Green cells include those matches where the query sequence and the corresponding database sequence have some overlap in the genome. Yellow cells have two numbers: the first number corresponds the total number of query sequences reported for each aligner, and the second number corresponds the number of reported query sequences that have a overlap with corresponding database sequence in the genome.

|  | *blastn* | *YASS* | *Water* | *Usearch* | *BWA-SW* | *Qudaich H I* | *Qudaich H II* |
| --- | --- | --- | --- | --- | --- | --- | --- |
| *blastn* | 9,999/<br>4,665 | 6,525 | 3,466 | 6,576 | 6,162 | 7,470 | 8,172 |
| *YASS* | 4,179 | 6,625/<br>4,182 | 3,354 | 6,428 | 6,127 | 5,932 | 6,116 |
| *Water* | 2,154 | 2,083 | 10,000/<br>2,430 | 3,310 | 3,213 | 3,170 | 3,242 |
| *Usearch* | 4,270 | 4,158 | 2,116 | 6,635/<br>4,292 | 6,147 | 6,028 | 6,210 |
| *BWA-SW* | 3,964 | 3,943 | 2,015 | 3,958 | 6,194/<br>4,176 | 5,684 | 5,842 |
| *Qudaich H I* | 4,387 | 3,950 | 2,057 | 4,052 | 3,772 | 10,000/<br>4,603 | 8,827 |
| *Qudaich H II* | 4,470 | 4,016 | 2,083 | 4,118 | 3,828 | 4,530 | 10,000/<br>4,625 |
| *Run time* | 1 min 25<br>sec | 28 min | >1 hour | 2.55 sec | 3.715 sec | 2.44 sec | 2.61 sec |

**Figure 5.**
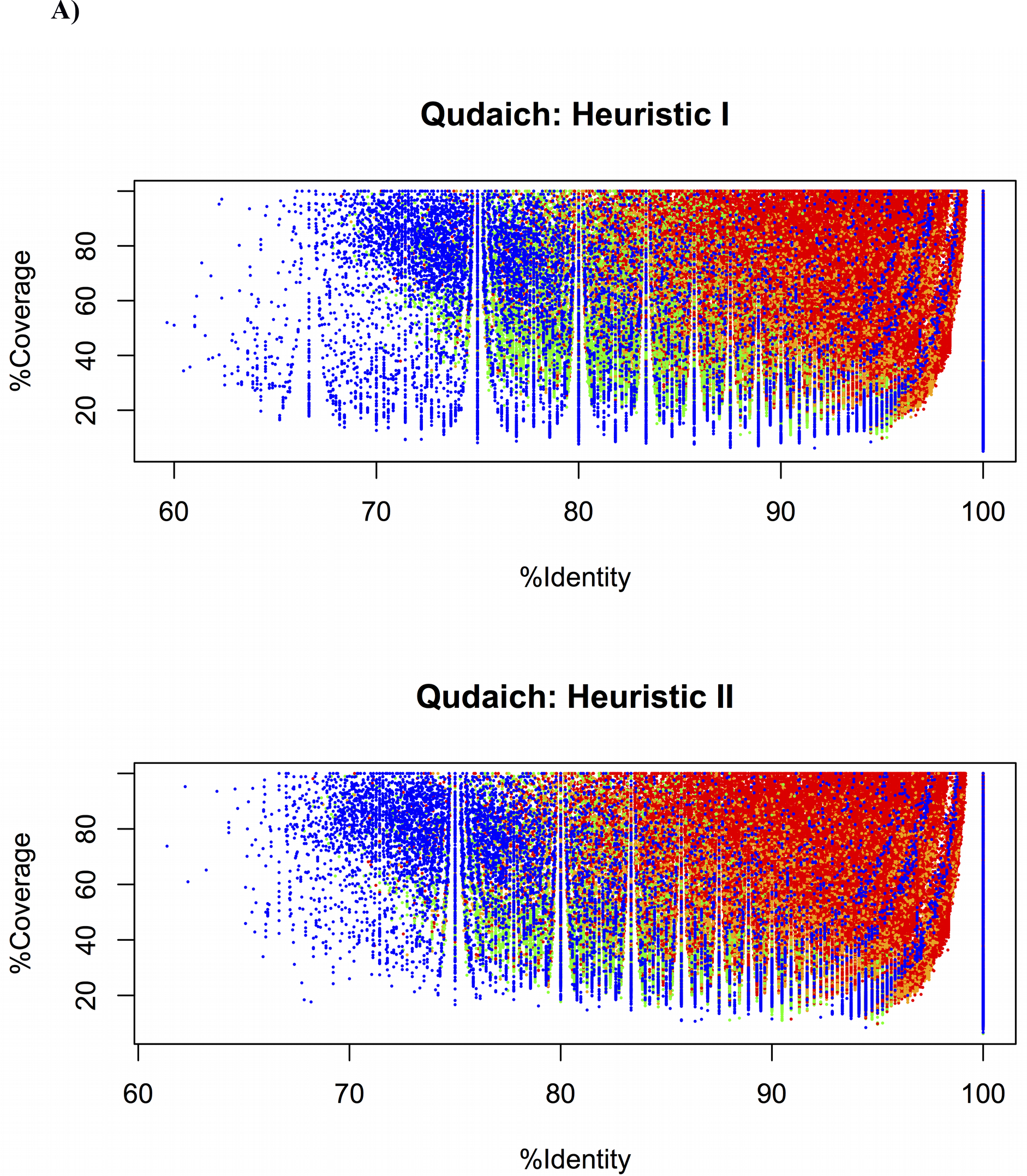
(a) The prediction of candidate database sequences. The top panel shows the prediction based on heuristic I, and the bottom panel shows the prediction based on heuristic II. The x-axis represents the % identity and y-axis shows the percent coverage of the alignment of each query sequence based on BLAST output. Different color shows the prediction of the candidate database sequence for each query sequence. Red and orange indicate those query sequences where the candidate database sequences identified by qudaich matches with the database sequences identified by BLASTn. The red color denotes those alignments that have the best score based on BLAST output. Blue indicates those queries where the database sequence predicted by BLAST does not match with the candidate database sequence predicted by qudaich. Green indicates those queries, which are reported in qudaich but not reported in BLAST output. (b) BLAST score based on the prediction of candidate database sequences. The color indicates the same group of candidate database sequences described in figure 4a.

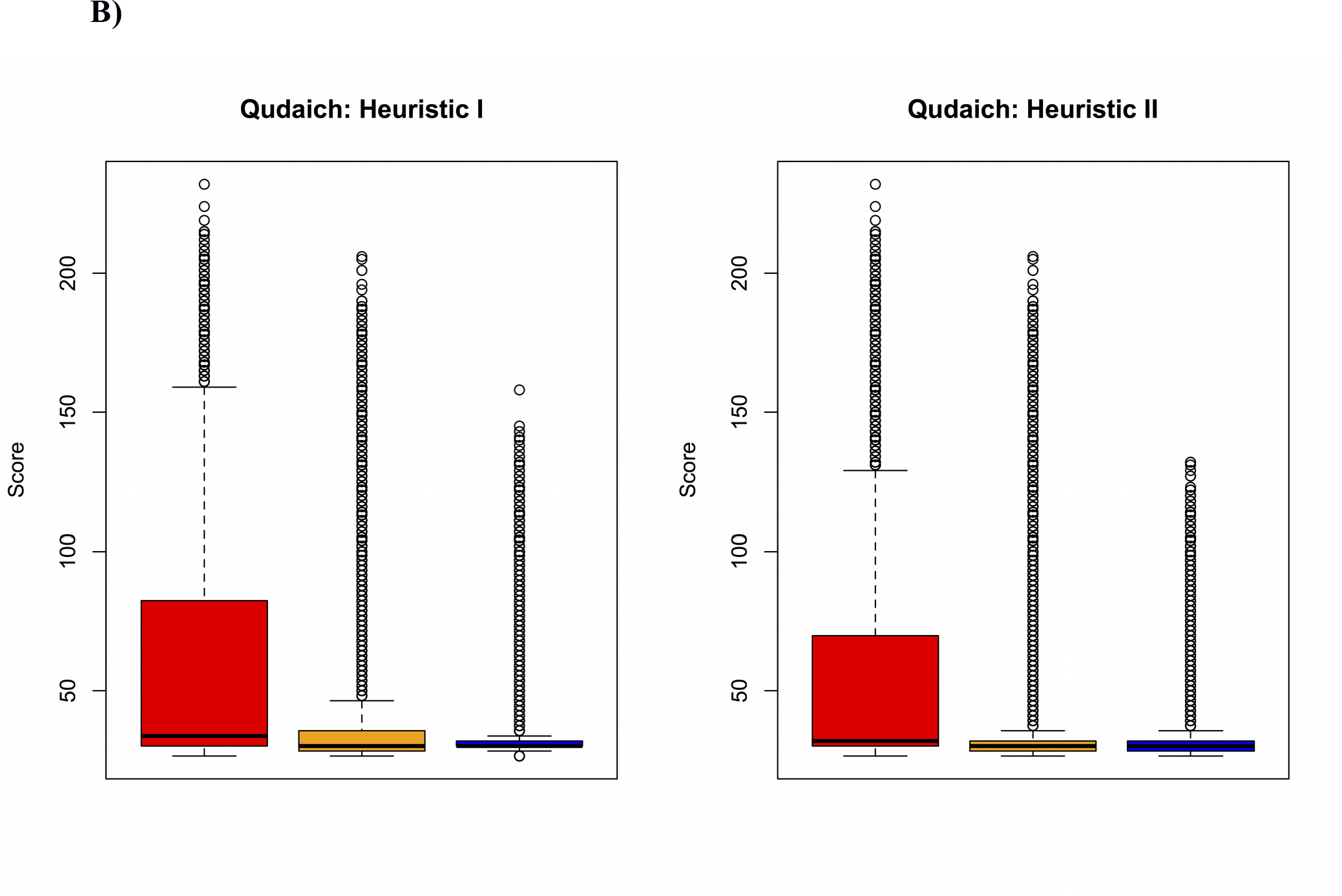

BLAST had the most matches alignments between query and database among all the algorithms tested (Table 1). Therefore, qudaich output was compared against the blastn output using a real NGS dataset (test set 2). Figure 5 shows the prediction of candidate database sequences based on the blastn results. When there is a good alignment between the query and the database sequences (high percent identity and high percent coverage), there is a strong correlation between qudaich predictions and those of blastn. When the coverage and identity are each greater than 90%, the candidate database sequence predictions match the blastn output 99.9% regardless of the heuristic used by qudaich (Table 2). The candidate database sequences that do not have a match with blastn output generally have poor alignment scores with low percent identity and low percent coverage (Figure 5a, Figure 5b). For these poor matches, using the top two database sequences per query sequence while calculating the candidate database sequence increases the similarity to the blastn output.

**Table 2:**
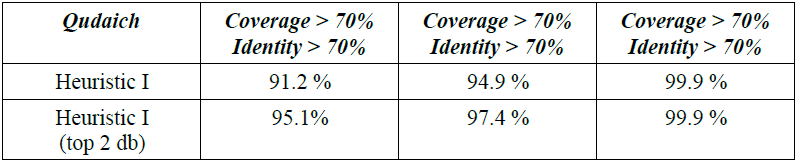
Percentage of predicted candidate database sequence matches with BLAST results using test set 1.

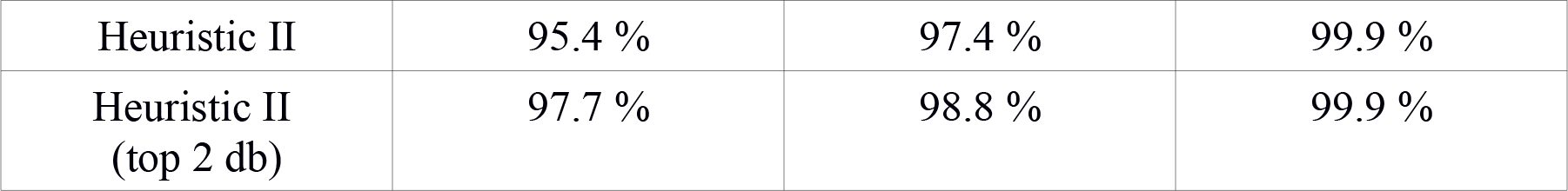

Testing the two different heuristics for selecting the database sequences showed that both approaches are effective at identifying the candidate database sequences. This suggested that the assertions underlying the two heuristics are validated by the data. However, when there is no strong match between the sequences, heuristic II gives better prediction than heuristic I, because of the inclusion of the lcp in the scoring. To investigate the reason why heuristic I gives poor performance for low % identity and low percent coverage, some alignments were manually checked. We explain this discrepancy using a hypothetical example. Suppose that for a given query sequence *q*, database sequence *d_1_* was chosen with heuristic I while database sequence *d_2_* was chosen with heuristic II, and let us assume that these are the only two sequences with homology to *q* in the database. When the alignments were examined between *q* and *d_2_*, we find a few longer exact matches. In contrast, when we examine the alignment between *q* and *d_1_* we find many shorter length matches. Heuristic I uses the frequency that the sequence occurs to identify the best match, so *d_1_* will dominate because of the many short matches. In contrast, heuristic II uses the total length of the longest common prefix to identify the candidate database sequence; therefore, sequence *d_2_* will dominate.

In the alignment result, if we consider only the good alignments (having high percent coverage and high percent identity) both heuristics actually find similar candidate
database sequences in this case. Therefore, qudaich can successfully predict the candidate database sequences.

### Real Time Statistics

A computer with an Intel(R) Xeon(R) CPU X3430@2.40GHz was used to compare and measure the execution time of qudaich and contemporary alignment algorithms. The run times of all programs would be enhanced by using multiple cores/threads (as discussed below), but for timing we used a single core/thread was used for all tests so that the results are comparable.

In qudaich, two different algorithms are designed for each of the two heuristics. Using test dataset 2, the time requirements for both algorithms were measured. The algorithm based on heuristic II required more time to construct suffix array and to find candidate database sequences because of the longest common prefix (lcp) calculation. As shown in Table 3, the SWG algorithm used to generate the alignments requires 73.5% and 66.6% of total execution time for heuristic I and heuristic II respectively. Hence, using a single instruction multiple data (SIMD) implementation of the SWG algorithm will probably reduce the total execution time of qudaich significantly.

**Table 3:** Real time statistics of different aligners for DNA sequence alignment using test set 2.

| *Aligner* | *Time* |
| --- | --- |
| blastn | 82 min 5 sec |
| YASS | 30 min 28 sec |
| WATER | > 1 day |
| USEARCH | 19 min 47.9 sec |
| Qudaich<br>Heuristic I | 2 min 14 sec<br>Construct SA: 22.2 sec<br>Find candidate DB sequences: 13.3 sec<br>Generate alignment: 98.5 sec |
| Qudaich<br>Heuristic II | 2 min 28.5 sec<br>Construct SA: 34.8 sec<br>Find candidate DB sequences: 14.8 sec<br>Generate alignment: 98.9 sec |

The alignment algorithms used above to compare the accuracy of qudaich output were also used in the timing for real time statistics. As shown in table 3, qudaich is about 33× faster than blastn, about 8× faster than USEARCH and about 12× faster than YASS.

### Protein sequence alignments

Unlike many other algorithms qudaich can also align protein sequences. For protein sequence alignments, qudaich was compared to the blastp program in BLAST+, RAPSearch, USEARCH and promer (integrated in MUMmer) using test dataset 3 (Kurtz et al., 2004; Edgar, 2010; Zhao, Tang & Ye, 2012). Table 4 shows that qudaich was faster than the other approaches for both protein vs. protein alignments and translated nucleotide vs. translated nucleotide alignments.

**Table 4:** Real time statistics for protein sequence alignment using test dataset 3.

| Aligner | Protein vs. protein | Translated nucleotide vs. translated nucleotide |
| --- | --- | --- |
| blastp or tbalstn | 124 min 2 sec | 757 min 43 sec |
| RAPSeach | 1 min 2.6 sec | 2 min 34.5 sec |
| USEARCH | 15.3 sec | - |
| PROMER | - | 4 min 48.9 sec |
| qudaich | 14.6 sec | 1 min 31.5 sec |

The qudaich output was compared with BLAST output for protein vs. protein sequence alignments (Figure 6). Like DNA sequence alignments, qudaich output agrees with BLAST output when there is an alignment with good percent coverage and good percent identity. However, for the alignments, where qudaich disagrees with BLAST output the alignment general has poor score (Figure 6).

**Figure 6.**
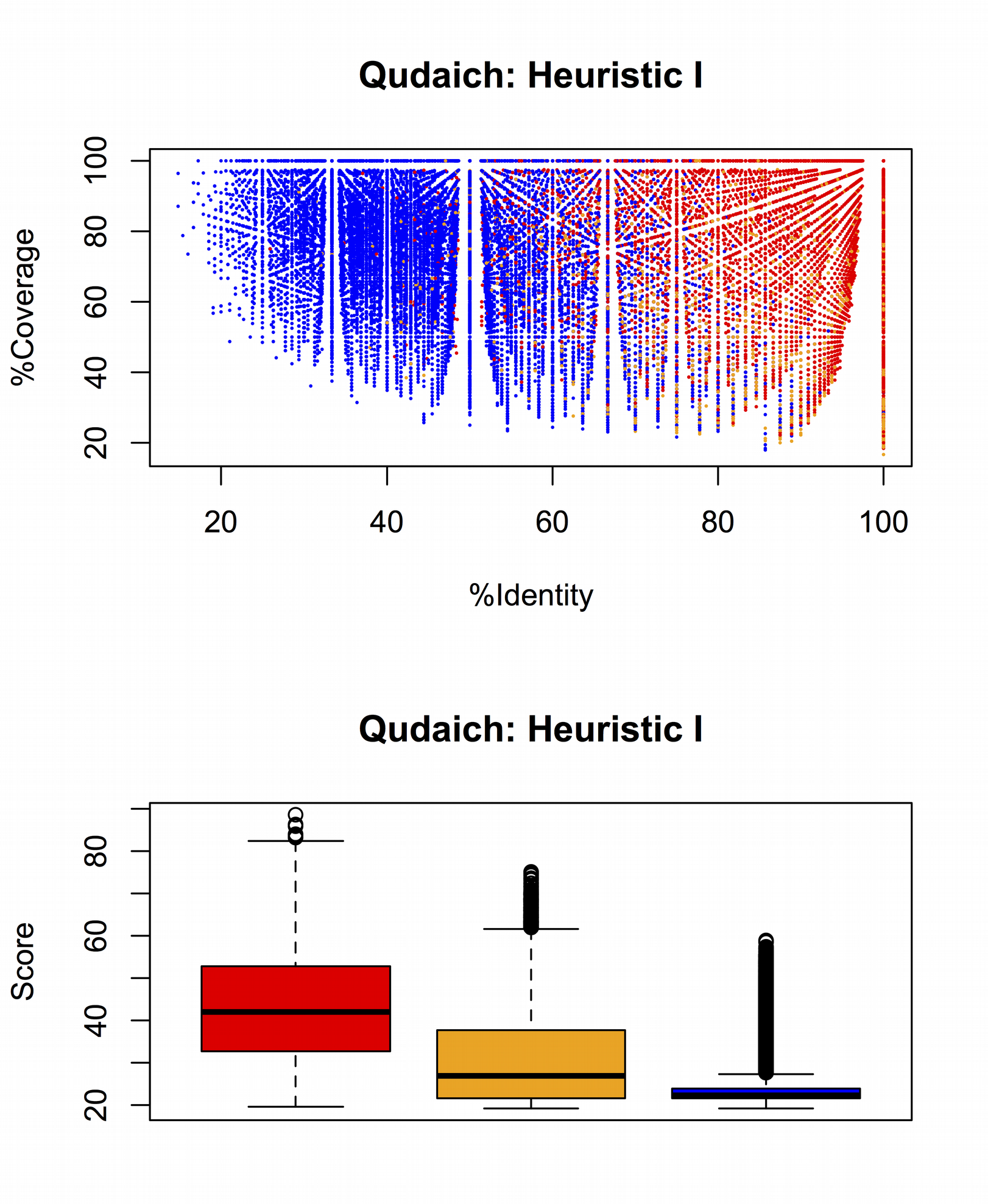
Prediction of candidate database sequences for protein sequence alignments using heuristic I. The top panel shows the prediction for each query sequences. The x-axis represents the percent identity and y-axis shows the percent coverage of the alignment of each query sequence based on blastp output. The bottom panel shows the blastp score based on the prediction of candidate database sequences. For both panel, the color indicates the same group of candidate database sequences described in figure 4a.

The above results show that qudaich produces more useful results for both DNA and protein sequence alignments but with less time than the contemporary approaches. This performance gain results from the novel algorithmic structure of qudaich that effectively reduces the search space for each query sequence to find the candidate database sequence. Qudaich is particularly suited to NGS datasets with huge numbers of database sequences.

## Discussion

Here we present a novel approach for local sequence alignment that tries to find the best possible alignment for each query sequence. In this approach, both database and query sequences are organized together. Based on this organization, the candidate database sequences are identified so that the optimal alignment can be generated using Smith-Waterman-Gotoh algorithm. In most of the cases, the alignments with the candidate database sequences are the best possible alignment against the whole database (Table 1, Figure 5). Two heuristics were developed for efficient searching of the candidate database sequences, and our algorithmic structure allows us to plug in different heuristics.

We showed that qudaich is more efficient in terms of execution time and accuracy than contemporary aligners. An interesting characteristic of qudaich is that, except for the suffix array construction, the time complexity mainly depends on the size of the query datasets. As the suffix array construction is linear, qudaich’s execution time should not be costly for large databases. This characteristic has a key advantage over the conventional approaches that find and extend or join seeds to construct alignments. In those cases, the execution time will be highly dependent on the database size. Aside from constructing the suffix array, the other steps of the algorithm can be easily parallelized. However, the only bottleneck of qudaich, comparing with the other aligners, is the memory usage for finding the candidate database sequences. Admittedly, qudaich constructs suffix array using both reference and query sequences, therefore it requires more memory than those approaches that rely on the Burrows-Wheeler algorithm or store only database sequences in memory.

With qudaich the percent similarity between two samples can be calculated without generating the actual alignments. In this case, qudaich gains a significant advantage over the other approaches in computation time: qudaich takes less than one third of total execution time (Table 3).

Our new sequence alignment algorithm, qudaich, will generate accurate sequence alignments between NGS datasets in less time, with less power, and less cost.

## Appendix

## Commands to run the aligners

## Qudaich v1

$ ./qudaich_search_db -query query.fasta -ref reference.fasta-freqFile FREQ.txt-prog n

$./qudaich_alignment -a all -freqFile FREQ.txt -output output.txt

## YASS v1.14

$ yass -d 2 -o output.txt query.fasta reference.fasta

## BLAST 2.2.29+

$ makeblastdb -in reference.fasta -dbtype nucl -out dbIndex

$ blastn -task blastn -db dbIndex -query query.fasta -out output.txt -outfmt ’6 std’

## Usearch v6.0.307

$ usearch6.0.307_i86linux32 -usearch_local query.fasta -db reference.fasta -strand both -threads 1 -evalue 1e-6 -id 0.5 -alnout output.txt

## Water

$ water -asequence query.fasta -bsequence db.fasta -gapopen 10.0 -gapextend 5 -outfile output.txt

## Promer (MUMmer3.23)

$ promer reference.fasta query.fasta

## RAPSearch v2.14

$ prerapsearch −d reference.fasta-n DB

$ rapsearch −q query.fasta -d DB -z 1 -v 1 -b 1

